# Structure-aware graph attention based hierarchical transformer framework for drug-target binding affinity prediction

**DOI:** 10.64898/2026.04.19.719524

**Authors:** Virendra Singh Kaira, Zakeerhussain D Kudari, Shankar P Sai, Ruchika Bhat, G. Jaiganesh

## Abstract

Drug-target interaction prediction is significant in the hit identification phase of drug discovery, enabling the identification of potential drug candidates for downstream optimization. Traditional computational methods have some drawbacks in their ability to represent 3D structural data for both molecules and target proteins, which is required for the intricate protein-ligand interactions that regulate binding affinity. In this approach, we propose a graph transformer-based model (GTStrDTI) that combines an intragraph attention mechanism with cross-modal attention to enrich the representation of both the drug molecule and target protein. This approach comprehensively models both intramolecular structural features and intermolecular interactions, thereby enhancing binding affinity prediction performance. A thorough evaluation on benchmark datasets such as KIBA, DAVIS, and BindingDB_Kd shows that our approach surpasses the state-of-the-art methods under challenging target cold-start settings. Our analysis found that augmenting graph-based 3D structural protein target (C-alpha contact graphs from PDB with threshold distance of 5Å) and incorporating molecule adjacency information, boosts predictive performance, thus contributing towards narrowing the gap between computational and experimental research.

## 1. INTRODUCTION

High-affinity binding between small molecules or peptides and their receptor proteins constitutes a fundamental selection criterion in the drug discovery process. Binding affinity can be measured experimentally [1] using techniques such as surface plasmon resonance, isothermal titration calorimetry, and radioligand binding assays. These methods are reliable but come with limitations: they are slow and expensive, especially when screening thousands of compounds at once. Consequently, the development of accurate computational models for binding affinity prediction [2] has become imperative to accelerate the drug discovery pipeline and reduce resource expenditure and time. Accurately predicting drug-target binding affinity offers critical insight into the pharmacological mechanisms. A systematic study of protein-ligand interaction is quite helpful in the hit identification phase. This is also beneficial in optimizing hit compounds, designing drugs, and identifying off-target effects.

Traditional methods for predicting drug target interactions can be categorized into three major categories: (i) ligand-based, (ii) structure-based and (iii) chemogenomic-based. Ligand-based focuses on quantitative structure-activity relationship (QSAR) models [3], which are governed by similarity principles, by using known bioactive molecules to infer potential interactions. It has limitations where the information about active ligand and sites is limited. Structure-based methods, such as molecular docking [4] and free energy perturbation (FEP) [5], leverage physics-based methods to simulate interactions at the atomic level. Being highly accurate, these methods, however, demand high computational resources and execution time. These methods are highly sensitive to force field and ligand parameters. The above-mentioned methods limit their ability in the virtual screening task. Chemogenomic [6] approaches represent a paradigm shift by employing machine learning algorithms to model drug-target relationships in a data-driven manner. These methods typically encompass three sequential stages:

i. data preprocessing and featurization of ligand molecular and protein structural data,
ii. training predictive models using diverse learning algorithms, and
iii. trained models used for affinity prediction on unseen drug-target pairs.

Our (GTStrDTI) method adopts a chemogenomic framework that leverages graph-based representations with a transformer [7] architecture for binding affinity prediction. This deep learning-based approach offers computational speed several orders of magnitude greater than physics-based simulations. However, its predictive performanceis critically dependent on the quality of molecular featurization and the architectural design of the underlying model. Despite these dependencies, the scalability and speed of machine learning methods position them as highly promising tools for large-scale virtual screening for hit identification.

## 2. RELATED WORK

Predicting how strongly a drug molecule binds to its target protein is one of the hardest problems in computational chemistry. Solving it could dramatically speed up the drug discovery process, from early screening to refining promising drug candidates. Molecular docking and virtual screening workflow employ a scoring function to rank candidate ligand poses within binding sites. Glide [8] and AutoDock Vina [9] are being extensively used as scoring functions. Simulation-based approaches such as free energy perturbation have also shown remarkable success in finding good binders. Their sensitivity to ligand parametrization and force field selection, combined with substantial computational demands, constrains their applicability to large compound libraries.

Machine learning has offered a faster and more practical alternative to traditional methods. One early example, RF-Score [10], took a straightforward approach; it examined how different types of atoms in the protein and drug molecules are arranged near the binding site, then used a random forest model to predict binding strength. Several methods that followed worked along similar lines with comparable results.

ID-Score [11], for instance, went a step further by pulling together 50 different descriptors that capture various aspects of how a drug interacts with its target, such as electrostatic forces, how well the two molecules fit each other’s shape, and how solvation affects the interaction. These descriptors were then fed into a Support Vector Machine to make predictions. In other studies, researchers combined descriptors from a popular docking tool called AutoDock Vina with random forest models, and managed to enhance accuracy even further. SFCscore (RF) [12] employed this strategy with the original AutoDock Vina descriptors and also shows similar results.

Deep learning techniques [13] are widely used in drug-target affinity (DTA) prediction. 1D CNNs are utilized in DeepDTA[14] to encode molecule SMILES and target protein sequence, followed by feature addition and classification modules. Graph Convolutional Networks (GCNs) are utilized in iEdgeDTA [15], where a 1D graph of protein sequence with edge attentive graph convolution of drug embeddings. Introduction of 3D convolutional neural networks (3D-CNNs) by KDEEP [16] outperformed alternative machine learning methods and classical scoring functions across multiple diverse datasets. To model drug-target interaction in a comprehensive manner, graph neural network (GNN) based architectures have been designed. GraphDTA [17] pioneered the application of GNNs for learning drug structural representations, integrating them with CNN-based protein sequence encoding. GraphPrint [18] constructs graph representations of protein three-dimensional structures using amino acid residue spatial coordinates, combining these with drug molecular graphs and traditional descriptors in a joint learning framework for DTA prediction. The generative AI model OmegaFold is used in DTA-GTOmega [19] for 3D structure generation of the target protein, and the graph encodes descriptor information. T-ALPHA [20] presents a hierarchical transformer-based model that enhances binding affinity prediction through integration of multimodal feature representation. AttentionMGT-DTA [21] introduces a multimodal attention-based architecture for DTA prediction, representing drugs as molecular graphs and targets as binding pocket graphs, incorporating dual attention mechanisms to facilitate information integration and interaction between protein modalities and drug-target pairs.

GTStrDTI methodology aims to mitigate the limitations of earlier studies by integrating advanced 3D structural information of both the target protein and the ligand through graph-based transformer networks since previous studies have been deficient in capturing the necessary features and architectures to realistically represent these interactions. By applying this approach, more sophisticated representations can be leveraged to achieve more accurate binding affinity predictions.

## 3. MATERIALS AND METHODS

### 3.1 Graph construction

#### 3.1.1 Drug Graph Representation

Drug SMILES (Simplified Molecular Input Line Entry System) are encoded into graph [22] features, which is shown in Figure 1, using RDKit [23]. Drug is represented by *G*_*d*_(*X*_*d*_, *E*_*d*,_ *E*_*xd*_), where 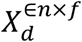 (n denotes the number of atoms, f are feature dimensions) represent the node features as listed in Table 1, 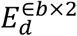 (b denotes the number of edges) represent the adjacency matrix, and 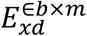 (m denotes the number of edge dimensions)

**Figure 1:**
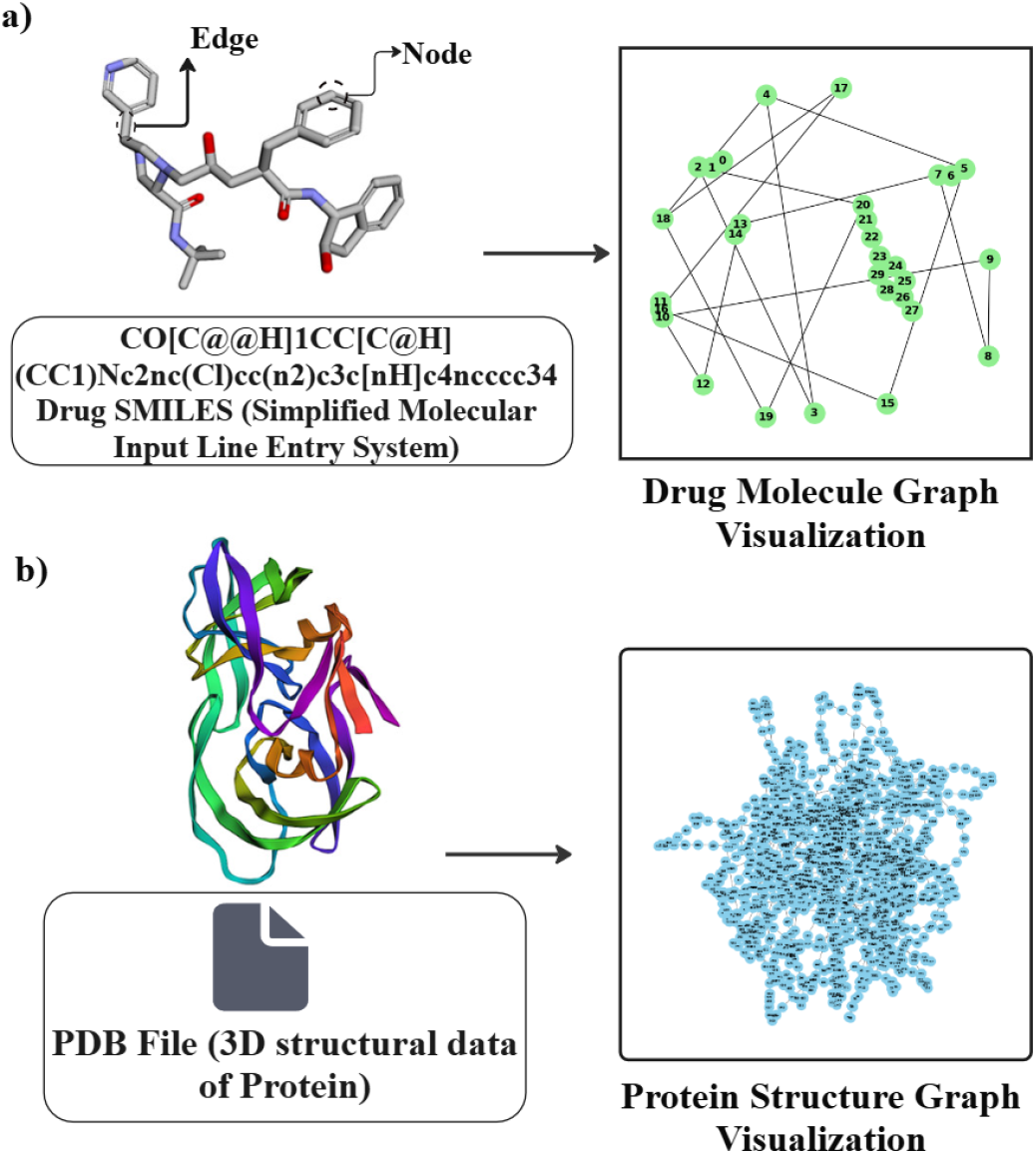
Graph-structured representation of **a)** drug molecules and **b)** protein targets, where atoms and residues form nodes and their structural relationships define edges.

**TABLE 1:**
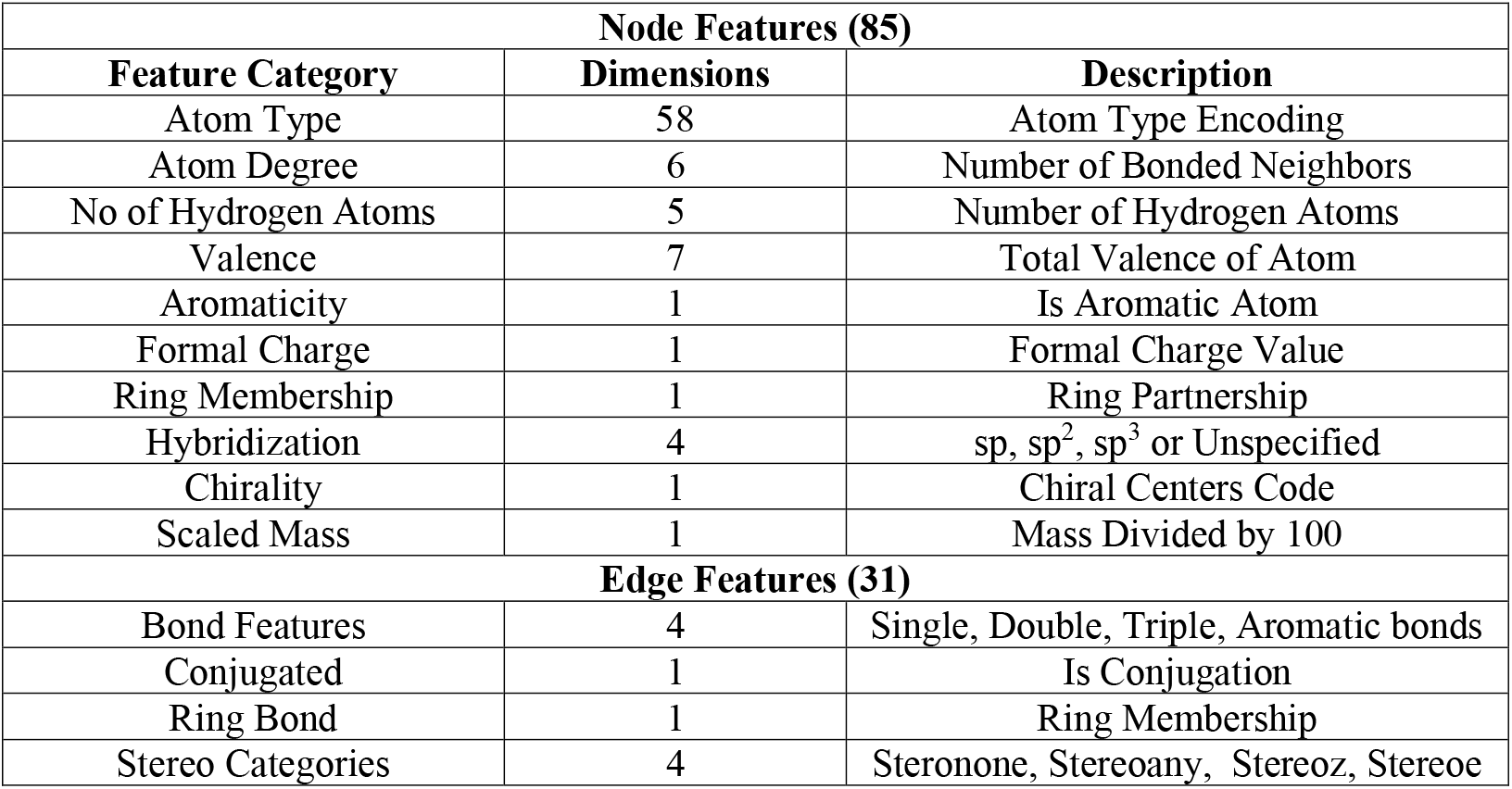
Feature Sets and Dimensional Specifications of Drug Molecule Graph Nodes and Edges.

| Node Features (85) |  |  |
| --- | --- | --- |
| Feature Category | Dimensions | Description |
| Atom Type | 58 | Atom Type Encoding |
| Atom Degree | 6 | Number of Bonded Neighbors |
| No of Hydrogen Atoms | 5 | Number of Hydrogen Atoms |
| Valence | 7 | Total Valence of Atom |
| Aromaticity | 1 | Is Aromatic Atom |
| Formal Charge | 1 | Formal Charge Value |
| Ring Membership | 1 | Ring Partnership |
| Hybridization | 4 | sp, sp<sup>2</sup>, sp<sup>3</sup> or Unspecified |
| Chirality | 1 | Chiral Centers Code |
| Scaled Mass | 1 | Mass Divided by 100 |
| Edge Features (31) |  |  |
| Bond Features | 4 | Single, Double, Triple, Aromatic bonds |
| Conjugated | 1 | Is Conjugation |
| Ring Bond | 1 | Ring Membership |
| Stereo Categories | 4 | Steronone, Stereoany, Stereoz, Stereoe |

#### 3.1.2 Protein Graph Representation

We encoded structural features into a graph representation by utilizing the *C*_*α*_ [24] centroid atomic centroids of each residue. Pairwise threshold distances of 5Å is applied to detect the edges. The protein graph is represented as *G*_*t*_(*X*_*t*_, *E*_*t*_), where 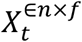 (n denotes the number of residues, f are feature dimensions) represent the node features, as illustrated in Table 2, 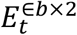 (b denotes the number of edges) represent the adjacency matrix.

**TABLE 2:** Integrated Node-Level Feature for Target Protein Graph Representation.

| Node Features (31) |  |  |
| --- | --- | --- |
| Feature Category | Dimension | Description |
| Amino Acid Type | 20 | 20 Standard Amino Acids |
| $C_{\alpha}$ centroid | 3 | $X, Y, Z$ Coordinates of $C_{\alpha}$ Atom |
| Hydrophobicity | 5 | Kyte-Doolittle Hydrophobicity Scale |
| Charge | 7 | Residue Charge at pH7 |
| Polarity | 1 | Binary Polarity |
| MW | 1 | Residue Molecular Weight (Dalton) |
| Aromaticity | 1 | Aromatic Ring Presence |
| H-Bond Donors | 4 | Number of Hydrogen Bond Donors |
| H-Bond Acceptors | 1 | Number of Hydrogen Bond Acceptors |
| Sidechain Volume | 1 | Volume of Amino Acid Sidechain |

### 3.2 Model

Figure 2 illustrates an advanced dual-stream transformer architecture that processes both drug and protein information as graph structures, representing a significant advancement in computational drug discovery. Our architecture processes drug and protein representations via parallel transformer pathways that share block configurations. It ensures symmetric feature extraction from both molecular entities prior to cross-modal integration via an attention mechanism.

**Figure 2:**
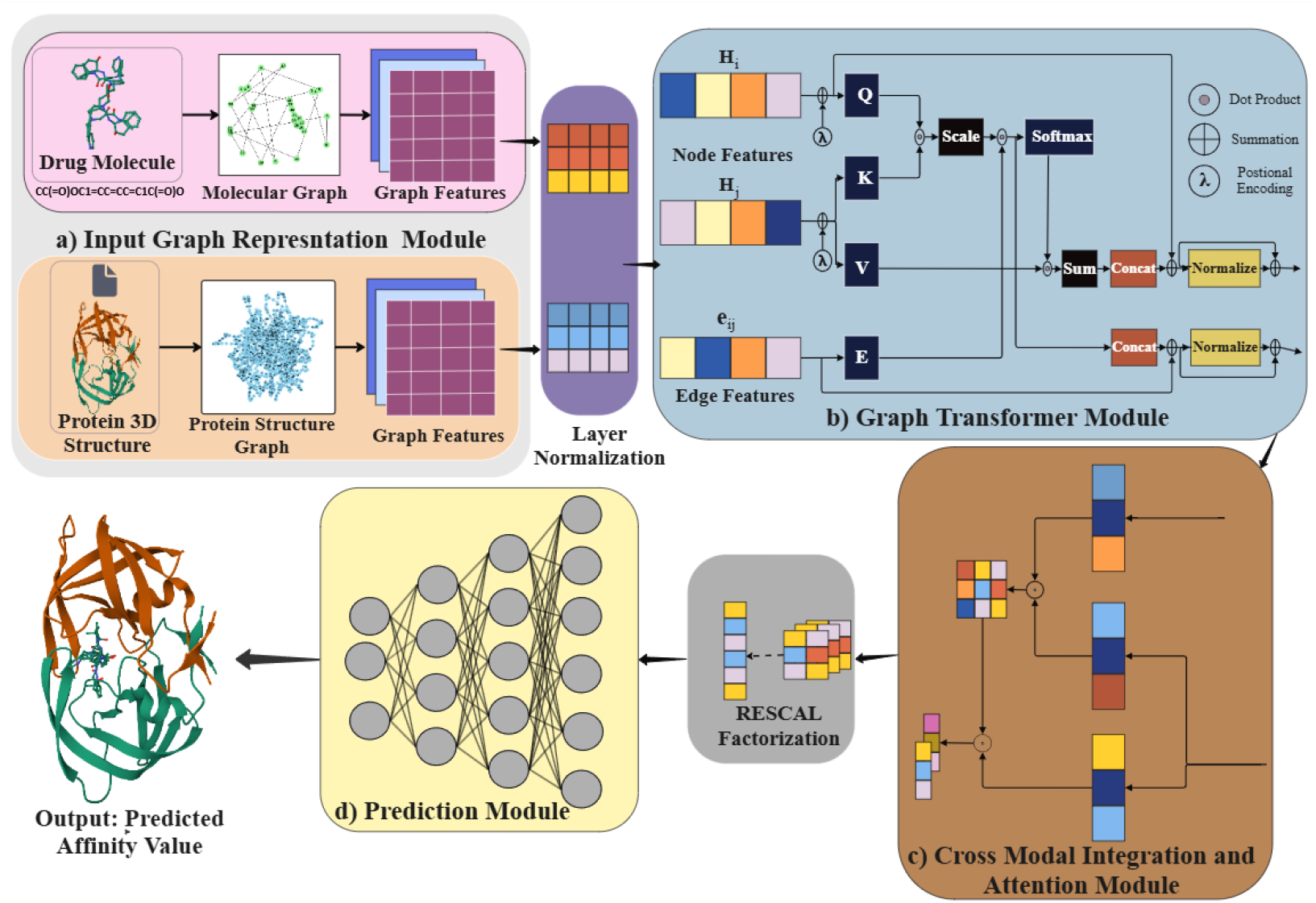
The framework consists of: **a)** Input Graph Representation Module: encodes molecules and proteins as graphs with atoms/residues as nodes and interactions as edge features; **b)** Graph Transformer Module captures local and long-range structural dependencies via transformer-based message passing; **c)** Cross-Modal Integration and Attention Module: fuses molecular and protein embedding using attention to model cross-modal interactions; **d)** Prediction Module uses the fused representation to predict interaction outcomes such as binding affinity or class labels.

#### 3.2.1 Transformer Block Architecture

The core innovation in our approach is the use of a dual transformer block, which takes a normalized graph representation as an input. Each transformer block has 4 essential modules arranged in a distinctive order:

##### a) Transformer Conv Layer

This specialized graph transformer layer extends traditional transformer attention mechanisms to graph structures. It computes attention weights between connected nodes, allowing the model to focus on chemically or structurally relevant regions. For drugs, this captures important pharmacophores [25] and reactive groups. For proteins, it identifies key binding residues and structural motifs.

##### b) Intra-Graph Attention

This component performs self-attention within each graph, enabling long-range dependencies and global context understanding. In molecular graphs, this allows atoms to attend to distant but chemically relevant atoms. In protein graphs, this captures allosteric effects and long-range structural constraints that influence binding.

##### c) SAG Pooling

Self-Attention Graph pooling provides a learnable method for aggregating node-level information into graph-level representations. Unlike traditional pooling methods that may lose important structural information, SAG [26] pooling maintains attention-weighted importance scores, preserving critical molecular or protein features while creating fixed-size representations suitable for downstream processing.

##### d) Graph Normalization + Exponential Linear Unit activation (ELU)

The combination of graph-specific normalization with ELU [27] ensures stable training dynamics while maintaining the expressiveness necessary for complex molecular and protein feature learning. ELU activation helps mitigate vanishing gradients common in deep graph networks.

The node features of both the drug and target graphs are subjected to Layer Normalization [28], which enhances training stability and accelerates the convergence of the neural network.

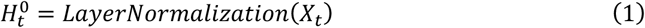

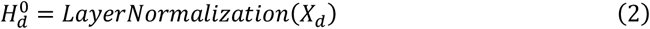

Now, normalized graphs 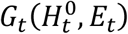, and 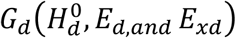 are passed to the graph transformer module, which applies the attention mechanism.

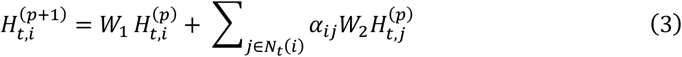

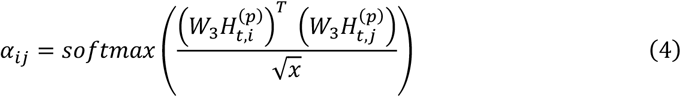

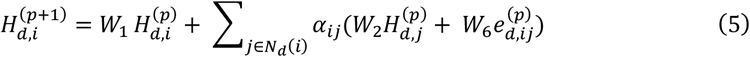

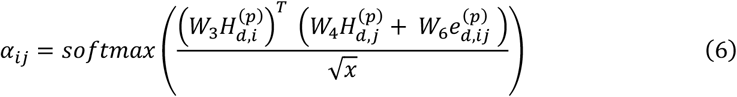

In this formulation, 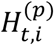 and 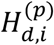 represent the feature vectors of node *i* at the *p*^*th*^ layer within the protein and drug graphs, respectively. The parameter *x* specifies the dimensionality allocated to each attention head in the hidden representation, while the matrices W correspond to trainable weights. Additionally, 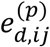 captures the edge features between nodes *i* and *j* at layer p in the drug graph *G*_*d*_, *and α*_*ij*_ is the attention weight showing how strongly node *i* attends to node *j*. After two layers of message aggregation, the updated node representations are obtained and denoted as *H*_*t*,(*p*)_ ∈ *R*^*n*×*v*^ and *H*_*d*,(*p*)_ ∈ *R*^*a*×*v*^, where m denotes the *p*^*th*^ Graph Encoder module. The dimension *v* corresponds to the product of the hidden layer size and the *x* number of attention heads. The outputs generated by each successive pair of Graph Transformer layers are subjected to the ELU activation function. Subsequently, SAG pooling and global pooling operations are employed to aggregate the node-level latent representations, yielding a compact one-dimensional global feature vector.

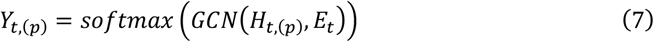

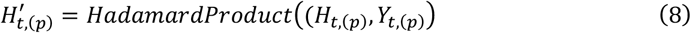

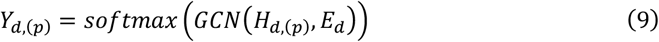

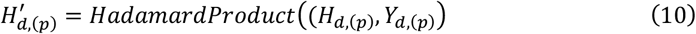

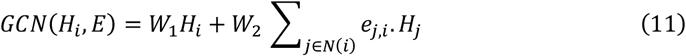

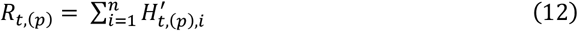

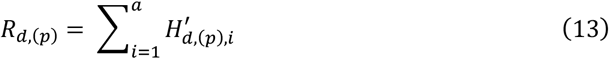

Here, the number of target residues is denoted by n, while the number of atoms in the drug molecule is denoted by a.

#### 3.2.2 Cross-Modal Integration and Attention

The architecture’s most sophisticated component is the Co-Attention [29] Layer, which facilitates interaction between processed drug and protein representations. This cross-modal attention mechanism allows the model to identify complementary features between drug and protein graphs, essentially learning which molecular features are most relevant for binding to specific protein regions. The co-attention mechanism computes attention weights between drug nodes and protein nodes, creating a rich interaction matrix that captures potential binding modes, steric complementarity, and electrostatic interactions. This approach goes beyond simple feature concatenation by learning dynamic, context-dependent interactions between molecular entities. After M serial Graph Encoder modules *G*_*t*_ and *G*_*d*_ are stacked to obtain *R*_*t*_ ∈ *R*^*M*×*V*^ and *R*_*d*_ ∈ *R*^*M*×*V*^, which are fed into the co-attention module to obtain pairwise interaction importance.

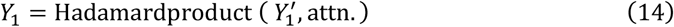

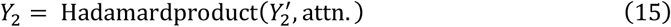

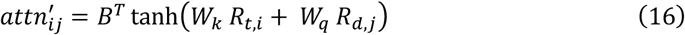

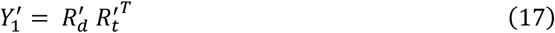

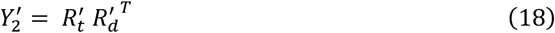

Here, 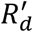 and 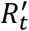 are the normalized forms of *R*_*t*_ and *R*_*d*_. *B*^*T*^, *W*_*k*_, and *W*_*q*_ are learnable matrices.

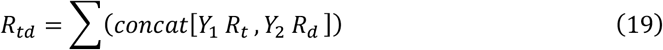

#### 3.2.3 Prediction Module

Following the co-attention integration, the architecture adopts a Rescale Tensor Factorization [30] module, which is designed to efficiently process the high-dimensional interaction tensors generated by the attention mechanism. This factorization approach reduces computational complexity while preserving essential interaction information necessary for accurate binding affinity prediction. The final representation (*R*_*td*_) of protein-drug interaction is passed to fully connected layers.

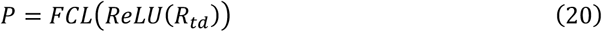

The final Output Layer transforms the factorized representations into binding predictions, typically regression values for binding affinity (*K*_*i*_).

## 4. EXPERIMENTS AND RESULTS

### 4.1 Dataset

#### 4.1.1 Description

The datasets in this work are curated for AI-driven virtual screening and binding affinity prediction tasks, including the KIBA dataset [31], Davis dataset [32], and BindingDB_*K*_*d*_ [33]. Each dataset integrates chemical and biological information necessary for ligand-based, structure-based, and hybrid QSAR (Quantitative Structure-Activity Relationship) modelling. We explored a range of open-source and proprietary datasets to understand their structure, quality, and suitability for predictive modeling, particularly for estimating binding affinities. This exploration is foundational to building robust QSAR models and screening millions of compounds in silico to discover potential hits for a given biological target. We have taken standard open-source datasets made available through institutional collaborations and licensed sources for training and validation purposes. The availability of diverse chemical and biological information in datasets made them substantial for our approach. The key fields included in the datasets are summarized below:

##### SMILES

It is a textual notation (encoding) used to represent chemical molecules in a linear string. The string “CC(=O)OCC” is a SMILES notation of ethyl acetate. In cheminformatics, it is predominantly used to represent molecules, compute descriptors, and encode input to various networks.

##### SMILES ID (Molecule or Ligand Identifier)

A unique identifier assigned to each molecule in the dataset, often derived from internal database IDs or external sources (e.g., BindingDB ID, PubChem CID). Helps in tracking, referencing, or mapping molecular data across different sources. E.g. “BDB123456” or “CID2244”

##### PDB ID (Protein Data Bank Identifier)

A unique 4-character alphanumeric code representing the 3D structure of a protein or protein-ligand complex in the Protein Data Bank. Enables access to detailed 3D structural information for docking and structure-based QSAR models. E.g., “1A2B” or “6LU7” (COVID-19 main protease). **Affinity Value** (Binding Affinity)

Quantitative measurement [34] of how strongly a ligand binds to its target protein, usually expressed in **IC**_**50**_, ***K***_***i***_, or ***K***_***d***_ units (typically in nanomolar, nM) used to rank compounds during virtual screening. It is the target variable for the model. Lower values indicate stronger affinity.

#### 4.1.2 Dataset Preprocessing

Remove Incomplete or Invalid Entries: SMILES, Protein Sequence, and Affinity Value. Standardize and validate SMILES, filter out invalid SMILES strings using RDKit (E.g.: molecules that cannot be parsed). Remove duplicates based on canonical SMILES.

Remove outlier: Remove outlier data based on affinity values and protein sequence length.

Standardize Affinity values: Convert Affinity Values to a Common Scale. Ensure all affinity values are in the same unit (E.g.: nM) and convert them into log *pK*_*d*_. Validate PDB IDs.

Target structure verification: Ensured PDB IDs are correctly formatted (4-character codes). We also removed any structures that were invalid or missing.

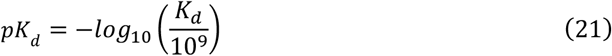

#### 4.1.3 Dataset EDA

Distribution plots of atom length, target sequence length, and datasets across all settings are provided in the Supplementary Information (SI) which is shown in Figures S1–S3, while the unique protein targets and drugs are summarized in Table 3. The Distribution of ligand atom length, target protein sequence length, and transformed activity values (p*K*_*d*_) indicates that the dataset includes small, medium, and large ligands interacting with protein targets of varying sequence lengths. The detailed Distribution is presented in Table 4.

**TABLE 3:** The table summarizes the dataset composition by reporting the total number of drugs– protein interaction pairs, together with the counts of unique drugs and unique proteins.

| Sl. No | Datasets | Unique Proteins | Unique Drugs | Total pairs |
| --- | --- | --- | --- | --- |
| 1 | KIBA | 1088 | 9887 | 42236 |
| 2 | DAVIS | 379 | 68 | 25772 |
| 3 | BindingDB Kd | 229 | 2086 | 117657 |

**TABLE 4:** Distribution of atom length characteristics in ligand SMILES, target Protein Length Range, Distribution of p*K*_*d*_ Values across Datasets.

| Distribution of Atom-Length |  |  |  |  |  |
| --- | --- | --- | --- | --- | --- |
| Sl. No | Datasets | Min | Max | Mean | Median |
| 1 | KIBA | 10 | 268 | 27.26 | 27.0 |
| 2 | DAVIS | 19 | 46 | 32.06 | 31.5 |
| 3 | BindingDB Kd | 5 | 389 | 33.70 | 32.0 |
| Distribution of Protein Sequence Length |  |  |  |  |  |
| 1 | KIBA | 215 | 4128 | 730.59 | 629 |
| 2 | DAVIS | 244 | 2549 | 744.85 | 632 |
| 3 | BindingDB Kd | 53 | 4128 | 725.75 | 629 |
| Range and Distribution of $pK_d$ | | | | | |
| 1 | KIBA | 7.76 | 8.96 | 7.93 | 7.94 |
| 2 | DAVIS | 5.0 | 10.79 | 5.41 | 5.0 |
| 3 | BindingDB Kd | 2.0 | 15.0 | 5.81 | 5.0 |

### 4.2 Experimental Settings

#### Training Configuration

The proportions of drug types in the training, validation, and test sets were set to 70%, 10%, and 20%, respectively, consistent with previously published studies to enable fair comparison. The detailed Distribution of the split datasets is provided in Table S1 of the Supplementary Material. Table 5 presents the experimental configuration used for model training.

**TABLE 5:** Experimental parameters used during training.

| **Parameters** |  |
| --- | --- |
| Batch Size | 32 |
| Learning Rate | 0.001 |
| epochs | 100 |
| patience | 50 |
| Optimizer | Adam |

#### Testing Configuration

In traditional evaluation methods, overlapping between training and testing data can lead to data leakage, resulting in models that perform exceptionally well during testing but poorly in real-world applications. The cold-start [35] setting, by strictly separating training and testing data, mitigates the risk of data leakage and overfitting. Furthermore, this setting can simulate real-world scenarios where new drugs or targets are encountered. By excluding these new drugs or targets during model training, we can assess the model’s generalization capability on unseen data, ensuring its effectiveness beyond just the training. Testing statistics are shown in Supplementary Information (SI) Table S1.

##### a. Drug cold-start setting

The training, validation, and test sets have only overlapped targets, but no overlapping drugs. For example, each test’s drug is new to the training and validation sets. The proportions of drug types in training, validation, and test sets are set to 70%, 10%, and 20%, respectively.

##### b. Target cold-start setting

The training, validation, and test sets have only overlapped drugs, but no overlapping targets. For example, each test’s target is new to the training and validation sets. The proportions of target types in training, validation, and test sets are set to 70%, 10%, and 20%, respectively.

##### c. Drug-target cold-start setting

The training, validation, and test sets have no overlap for both drugs and targets. For example, each test’s drug and target are both new to the training and validation sets. The proportions of drug and target types in training, test, and validation sets are set to 70%, 20%, and 10%, respectively.

### 4.3 Evaluation Metrics

We assessed the performance of the DTA prediction models using commonly used regression metrics, including Mean Squared Error (MSE), Root Mean Square Error (RMSE), Spearman rank correlation coefficient (ρ), Pearson correlation coefficient (r), and the concordance index [36] (CI). Lower RMSE and MSE values indicate smaller deviations between predicted and observed DTA values, whereas higher values for the remaining metrics indicate stronger agreement between model predictions and ground-truth DTA/DTI outcomes.

#### a) Root Mean Square Error (RMSE)

Root Mean Squared Error (RMSE) evaluates the average magnitude of prediction errors, giving higher importance to larger errors, and is expressed in the same units as the target variable.

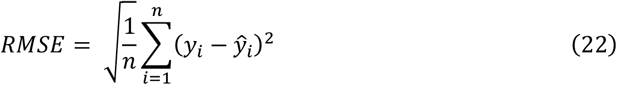

Where n= number of observations, *y*_*i*_ = actual value, *ŷ*_*i*_ = predicted value

#### b) Mean Squared Error (MSE)

MSE quantifies prediction error as the mean of the squared deviation between actual and predicted values. It is the foundation of RMSE.

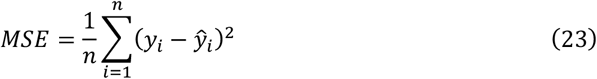

Where n= number of observations, *y*_*i*_ = actual value, *ŷ*_*i*_ = predicted value

#### c) Pearson Correlation Coefficient (r)

Provides a quantitative measure of the linear association between two continuous variables. It is the parametric test assuming normal distributions. Values range from −1 to +1. r = 1 represents perfect positive linear relationship, r = −1 perfect negative linear relationship and r = 0 shows no linear relationship.

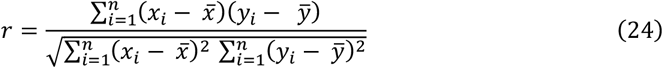

Where n = number of observations, *x*_*i*_, *y*_*i*_ = individual data points, 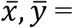 means of x and y

#### d) Spearman Rank Correlation Coefficient (ρ)

It represents monotonic correlation based on the ranks of data values rather than the raw values. It measures a monotonic relationship. Values range from −1 to +1. p = 1 represents perfect positive linear relationship, p = −1 perfect negative linear relationship and p = 0 shows no linear relationship. It is useful for non-normal distributions when we want to assess a monotonic rather than a linear relationship.

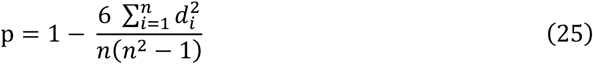

Where n = number of observations, *d*_*i*_ = difference between ranks of paired observations

#### e) Concordance Index (CI)

Concordance Index (CI) measures how well a predictive model preserves the correct ranking of outcomes. It calculates the proportion of all valid sample pairs for which the predicted values are ordered correctly, giving full credit for correct order (1), half credit for ties (0.5), and no credit for incorrect order (0). CI ranges from 0 to 1, with higher values indicating better ranking performance.

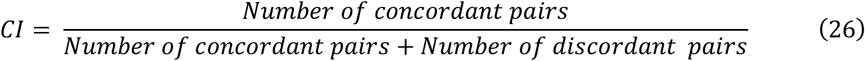

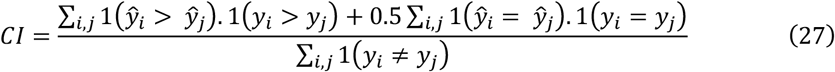

### 4.4 Results

We compare the proposed method with existing mainstream DTA prediction models across three benchmarks: drug cold-start, target cold-start, and drug-target cold-start settings. Scatter plots for all settings are in Supplementary Information (SI) Figures S4-S6. We have achieved a significant improvement over our previous counterpart in target cold start testing, highlighted in Table 6. The target cold-start setting evaluates a model’s ability to predict interactions for previously unseen protein targets. Most therapeutic proteins have limited or no experimental interaction data. A model that performs well only on targets it has already seen is not useful for novel target exploration. It also reduces overestimation of performance. Warm-start or random splits can inflate accuracy because the model indirectly sees similar versions of the same protein in training.

**TABLE 6:** Comparative Performance Analysis of Multiple Models under Drug Cold-Start Conditions on the DAVIS Dataset, KIBA, and BindingDB Datasets.

| Metrics | GTStrDTI<br>(Ours) | GraphDTA<br>(GAT_GCN)* | GraphDTA<br>(GIN)* | DrugMGR* | DTA_GTOmega* |
| --- | --- | --- | --- | --- | --- |
| **DAVIS Dataset** |  |  |  |  |  |
| **Drug cold start** |  |  |  |  |  |
| RMSE | 0.7863 | 0.755 | 0.733 | 0.970 | 0.692 |
| MSE | 0.6183 | 0.570 | 0.537 | 0.940 | 0.479 |
| Pearson (r) | 0.2592 | 0.322 | 0.233 | 0.248 | 0.424 |
| Spearman (q) | 0.2250 | 0.240 | 0.189 | 0.260 | 0.382 |
| CI | 0.6385 | 0.649 | 0.617 | 0.661 | 0.737 |
| **Target cold start** |  |  |  |  |  |
| RMSE | **0.7029** | 0.756 | 0.773 | 0.942 | 0.746 |
| MSE | **0.4940** | 0.571 | 0.597 | 0.888 | 0.556 |
| Pearson (r) | **0.5370** | 0.455 | 0.451 | 0.433 | 0.457 |
| Spearman (q) | **0.4590** | 0.432 | 0.360 | 0.360 | 0.440 |
| CI | **0.7532** | 0.737 | 0.698 | 0.698 | 0.742 |
| **Drug Target cold start** |  |  |  |  |  |
| RMSE | 0.8132 | 0.746 | 0.790 | 1.329 | 0.722 |
| MSE | 0.6613 | 0.557 | 0.625 | 1.767 | 0.522 |
| Pearson (r) | 0.0640 | 0.068 | 0.031 | -0.146 | 0.160 |
| Spearman (q) | 0.0644 | 0.105 | 0.011 | -0.092 | 0.203 |
| CI | 0.5409 | 0.567 | 0.507 | 0.441 | 0.628 |
| **KIBA Dataset** |  |  |  |  |  |
| **Drug cold start** |  |  |  |  |  |
| RMSE | 0.6586 | 0.659 | 0.635 | 0.752 | 0.629 |
| MSE | 0.4338 | 0.434 | 0.403 | 0.565 | 0.396 |
| Pearson (r) | 0.6478 | 0.64 | 0.668 | 0.485 | 0.689 |
| Spearman (q) | 0.6219 | 0.637 | 0.664 | 0.449 | 0.669 |
| CI | 0.7395 | 0.747 | 0.761 | 0.669 | 0.763 |
| **Target cold start** |  |  |  |  |  |
| RMSE | 0.6800 | 0.692 | 0.712 | 0.709 | 0.657 |
| MSE | 0.4624 | 0.478 | 0.507 | 0.503 | 0.432 |
| Pearson (r) | 0.5824 | 0.575 | 0.526 | 0.535 | 0.624 |
| Spearman (q) | **0.5117** | 0.504 | 0.429 | 0.489 | 0.507 |
| CI | **0.692** | 0.691 | 0.660 | 0.686 | 0.692 |
| **Drug Target cold start** |  |  |  |  |  |
| RMSE | 0.8103 | 0.842 | 0.838 | 2.667 | 0.787 |
| MSE | 0.6566 | 0.708 | 0.703 | 7.111 | 0.619 |
| Pearson (r) | 0.4004 | 0.365 | 0.337 | 0.105 | 0.438 |
| Spearman (q) | **0.4207** | 0.321 | 0.307 | 0.123 | 0.368 |
| CI | **0.6539** | 0.616 | 0.610 | 0.544 | 0.634 |

| BindingDB dataset |  |  |  |  |  |
| --- | --- | --- | --- | --- | --- |
| **Drug cold start** |  |  |  |  |  |
| RMSE | 1.1210 | 1.080 | 1.419 | 0.979 | 0.994 |
| MSE | 1.2566 | 1.166 | 2.013 | 0.958 | 0.988 |
| Pearson (r) | 0.6428 | 0.678 | 0.276 | 0.732 | 0.725 |
| Spearman (q) | 0.5840 | 0.621 | 0.217 | 0.670 | 0.681 |
| CI | 0.7392 | 0.758 | 0.585 | 0.778 | 0.784 |
| **Target cold start** |  |  |  |  |  |
| RMSE | 1.2553 | 1.279 | 1.238 | 1.233 | 1.214 |
| MSE | 1.5759 | 1.637 | 1.532 | 1.519 | 1.474 |
| Pearson (r) | 0.4488 | 0.379 | 0.459 | 0.451 | 0.465 |
| Spearman (q) | **0.4341** | 0.370 | 0.427 | 0.400 | 0.421 |
| CI | **0.6869** | 0.653 | 0.678 | 0.669 | 0.675 |
| **Drug Target cold start** |  |  |  |  |  |
| RMSE | 1.3778 | 1.428 | 1.386 | 1.363 | 1.324 |
| MSE | 1.8984 | 2.038 | 1.921 | 1.859 | 1.753 |
| Pearson (r) | 0.2773 | 0.206 | 0.353 | 0.273 | 0.386 |
| Spearman (q) | 0.2569 | 0.180 | 0.312 | 0.237 | 0.353 |
| CI | 0.6010 | 0.571 | 0.624 | 0.595 | 0.640 |

\* Reference [19]

### 4.5 Case Studies

We validate the practical readiness of approach through extensive case studies spanning diseases across four major therapeutic categories: respiratory, cardiovascular, infectious, and neurological disorders. These diseases are chosen so that the holistic efficacy of the model can be tested across different types of protein targets and verify the generalizability of the model. Results are shown in Table 7, which involves the comparison of predicted binding affinity of inhibitors for the unseen crystal structure of HIV-1 Protease (PDB ID: 1JKH, 6DH3, 3S56, 2AQU, 8FN7, 4MBS, 2ZD1, 1MUI, 2B60, 6PUW), where average standard deviation between experimental and predicted binding affinity values is found to be 1.29 Kcal/mol. Comparison results that can be referenced from table were also performed on crystal structures of Human VMAT2 associated with Huntington’s disease (PDB IDs: 8T6A, 8JTA, 3WS9, 3JSI, 5W5V, 4PH9) target protein with inhibitors, shows standard deviation in binding affinity values against experimental results is 0.97 Kcal/mol. We have compared inhibitor binding affinity for crystal structures of Non-Small Cell Lung Cancer (NSCLC) (PDB IDs: 7JXM, 2XP2, 4XV2, 7JUR) correspond to experimental data, results show a standard deviation of 0.87 kcal/mol. Unseen crystal structure of a congenital heart disease-associated protein (PDB IDs: 7DDH, 2H42, 2C6N) target with its some inhibitor was selected for comparison and results show a standard deviation of 0.73 kcal/mol.

**Table 7:**
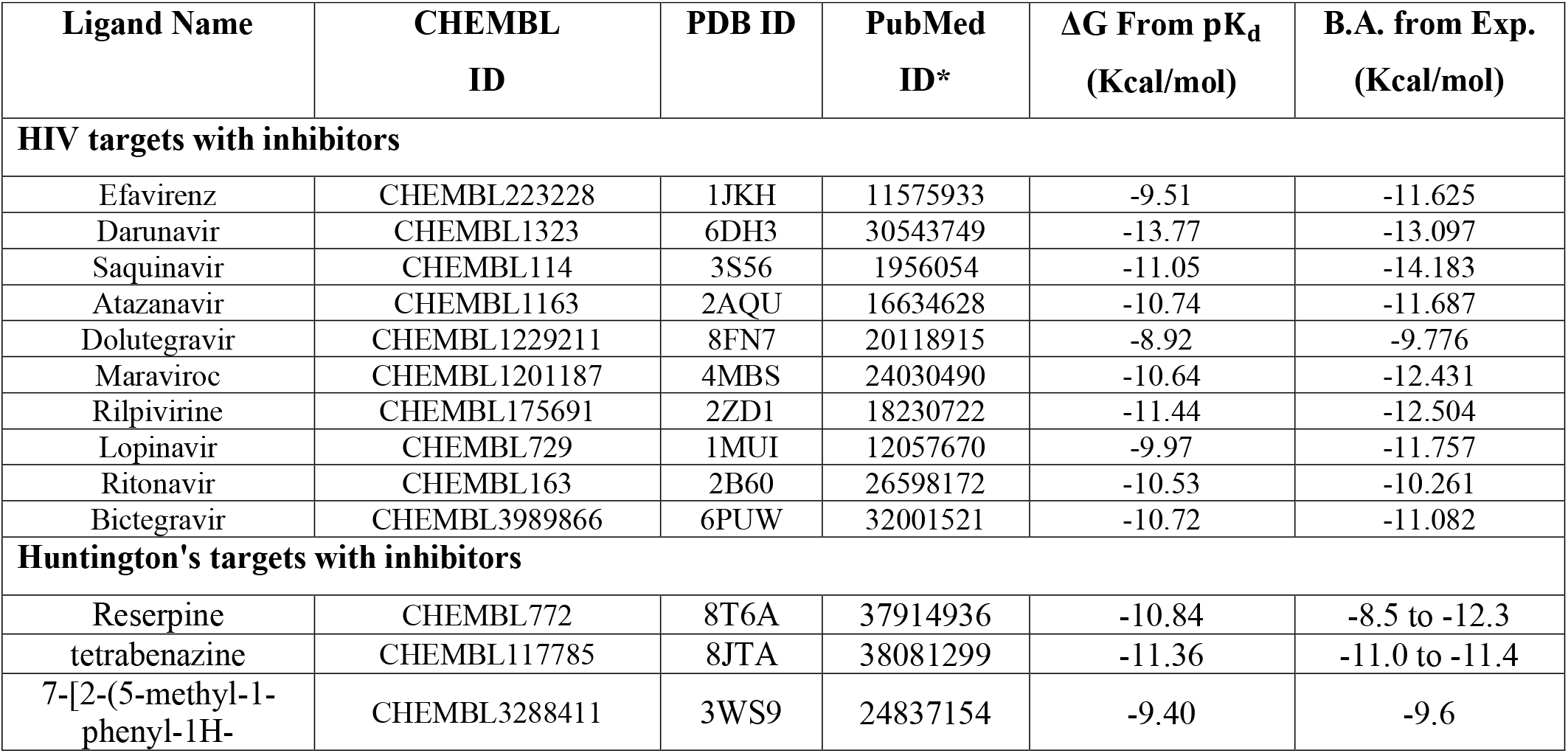
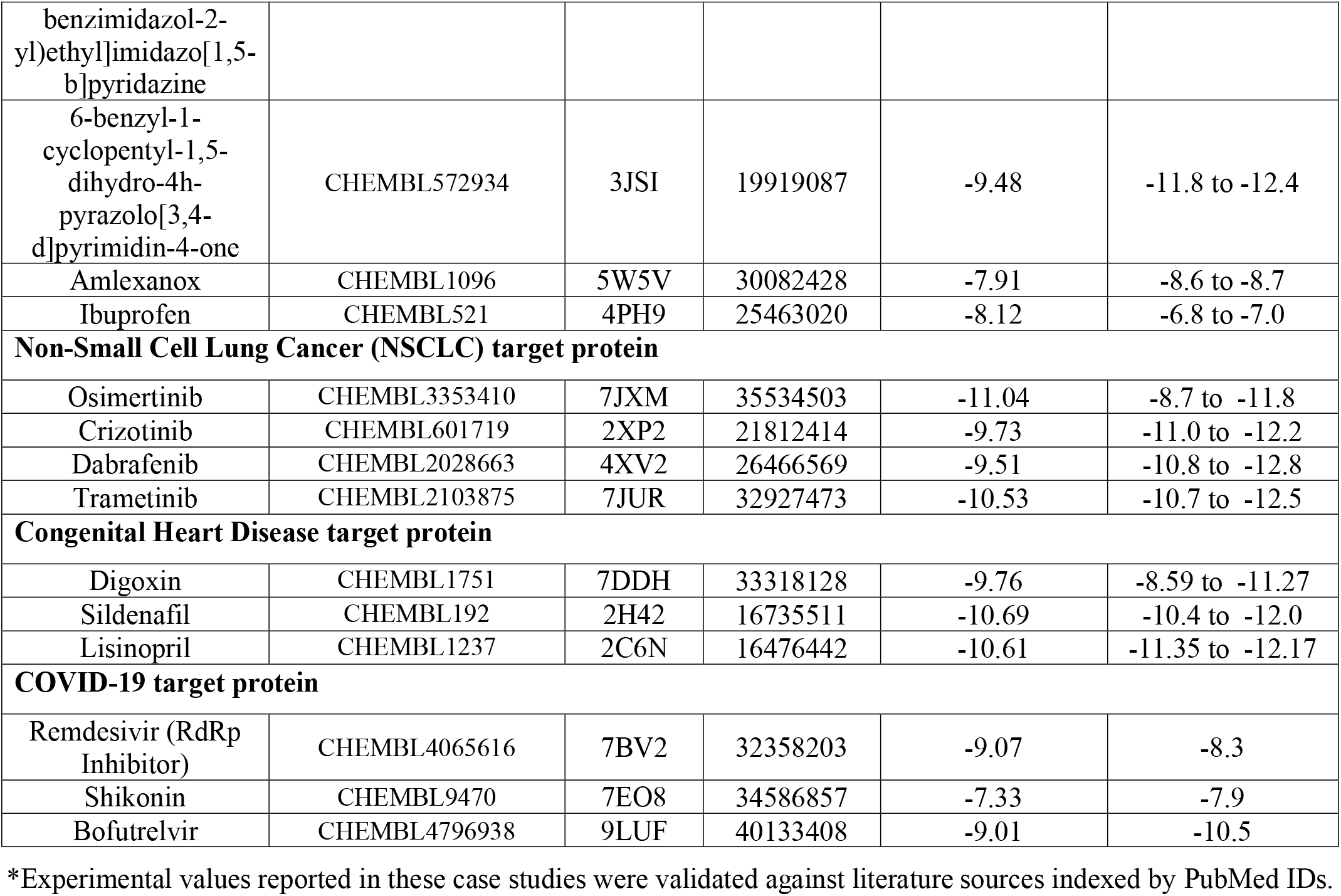
Comparison of GTStrDTI predictions with experimentally reported values for inhibitor molecules targeting the HIV-1 protein, Huntington’s targets, Non-Small Cell Lung Cancer (NSCLC) target, congenital heart disease target protein, highlighting agreement between computational predictions and experimental results.

As emphasized in the Table 7, the 3D crystal structure of the SARS coronavirus main protease (PDB IDs: 7BV2, 7EO8, 9LUF) target taken with inhibitors to infer the prediction results and the deviation from experimental results, corresponds to a standard deviation of 0.94 kcal/mol, showed strong alignment and correlation, indicating the robustness of the proposed approach for varied molecule target pairs.

## 5. CONCLUSION

We presented a framework for predicting drug target binding affinity that integrates 3D structural, physicochemical, electronic, and bond properties of both drug molecules and protein targets. It can operate without the need for highly computational pretrained models for feature extraction or prediction. This approach is flexible in processing artificially generated and experimentally determined protein targets. We have demonstrated our model’s capability to enhance drug candidate selection during virtual screening, thereby accelerating the drug discovery pipeline by reducing both time and cost requirements. Through the integration of 3D protein structural information, our model successfully addresses the challenging target cold-start scenario across the DAVIS, KIBA, and BDB datasets, outperforming existing methods that depend exclusively on conventional protein representations or pretrained models. Given the recent advances in accurate computational protein structure prediction and the expanding availability of 3D protein structure databases, incorporating structural information represents a promising new paradigm for drug-target affinity prediction and related applications in drug discovery and multimodal learning. This research improves our understanding of drug target interaction. It provides practical benefits by helping make the drug development process more efficient and cost-effective. These evaluation studies demonstratethe model’s utility in computational drug discovery.

However, the approach has some limitations. Notably, our work does not include an ablation study, making it difficult to evaluate each component’s contribution. Additionally, limited explainability of deep learning is one of the hindrances in trusted adoption of the technique. With further improvements in features representation by considering secondary structure, torsion Angle and dynamics features, it could become even more accurate in its predictions. On the architecture side, adapting to optimize graph pooling, sparse attention and tweaking the positional encoding can further enable researchers to identify promising drug candidates with improved computational efficiency.

## ACKNOWLEDGEMENTS

The authors would like to express their sincere gratitude to all individuals who contributed their time, expertise, and support during the course of this research.

## DECLARATIONS

The authors declare that they have no competing interests.

## Supplementary Information

**Figure S1:**
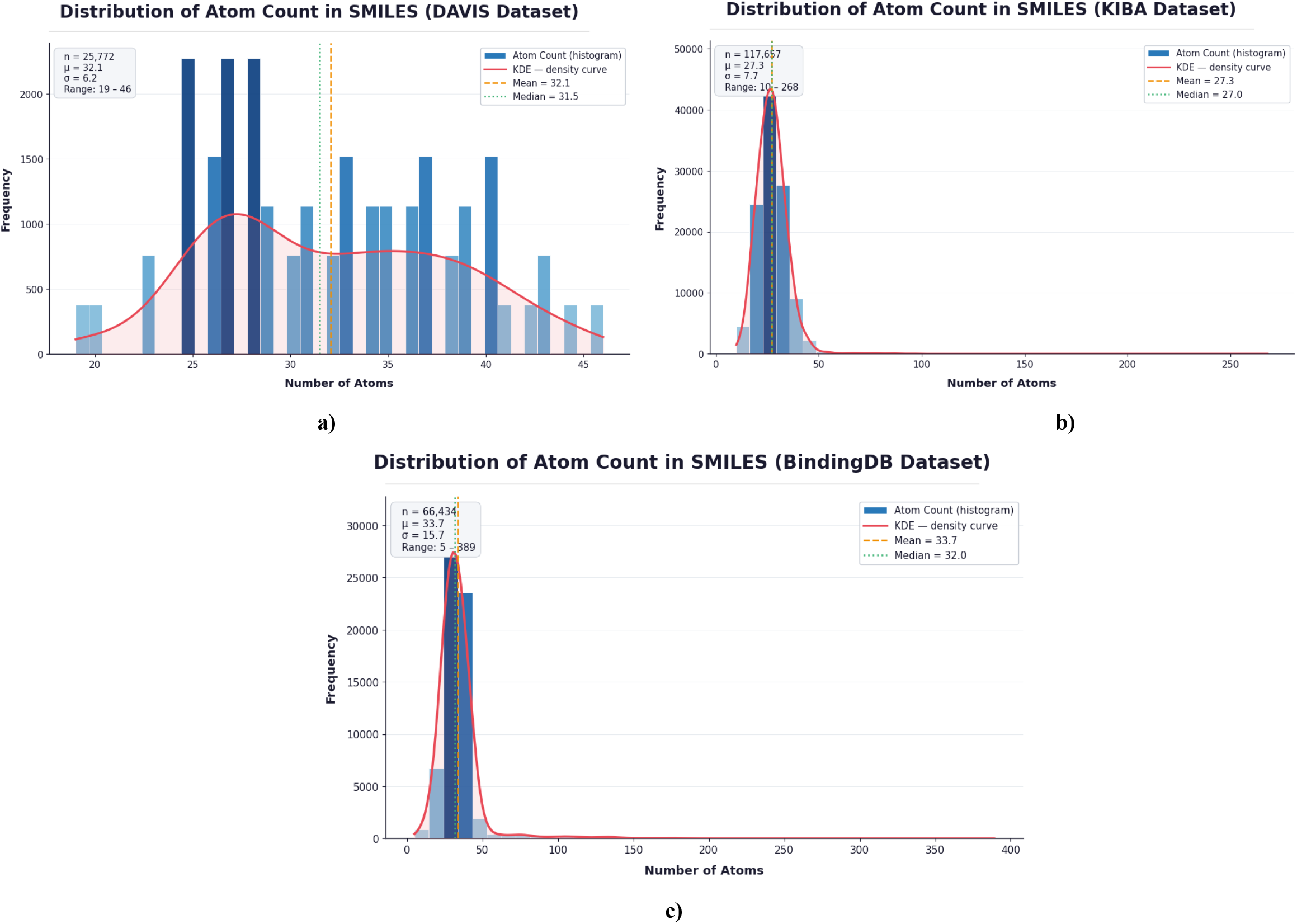
Distribution of Molecule (SMILES) atom count; a) DAVIS Dataset, b) KIBA Dataset, c) BDB Dataset

**Figure S2:**
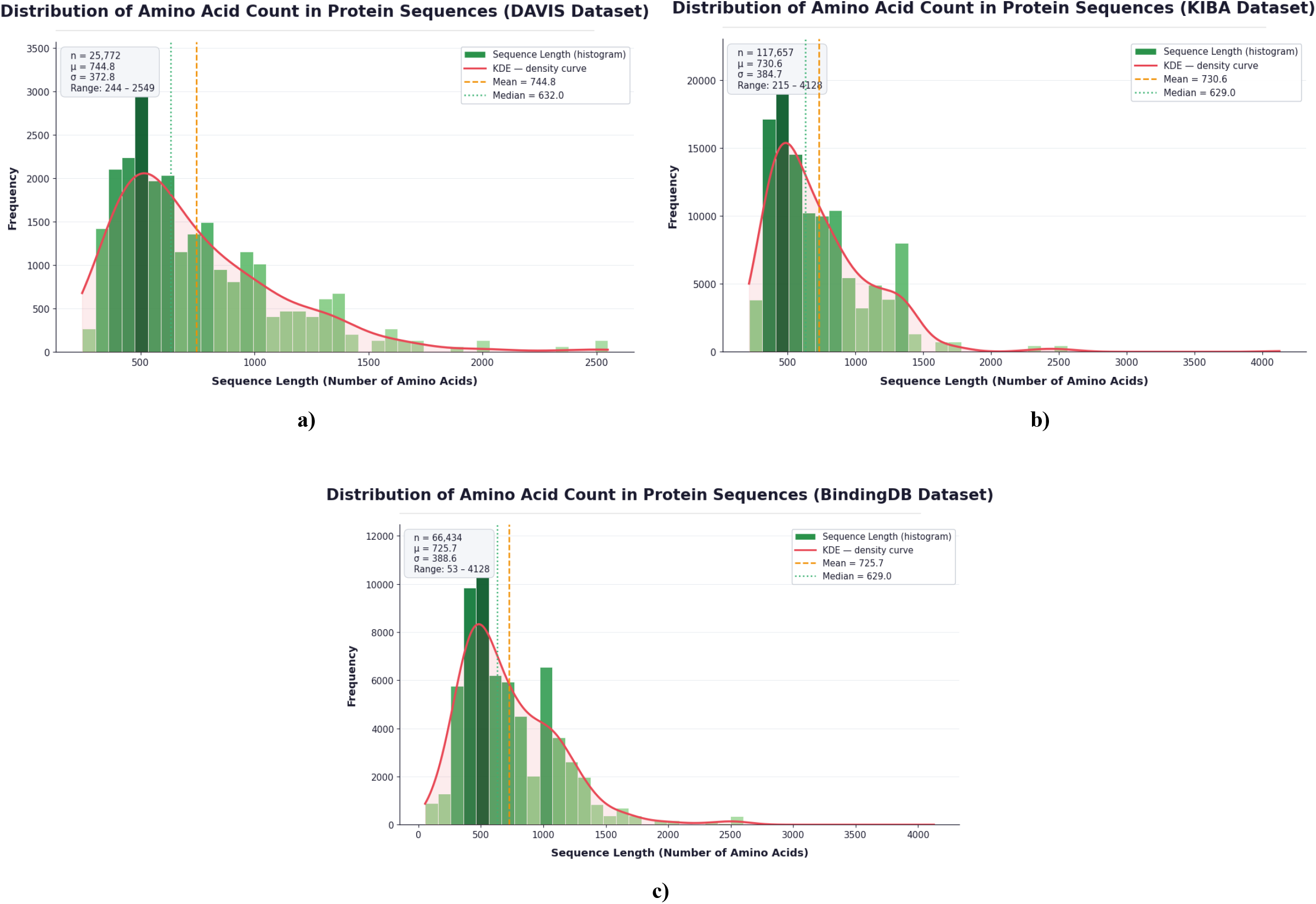
Distribution of amino acid count in protein sequence of : a) DAVIS Dataset, b) KIBA Dataset, c) BDB Dataset

**Figure S3:**
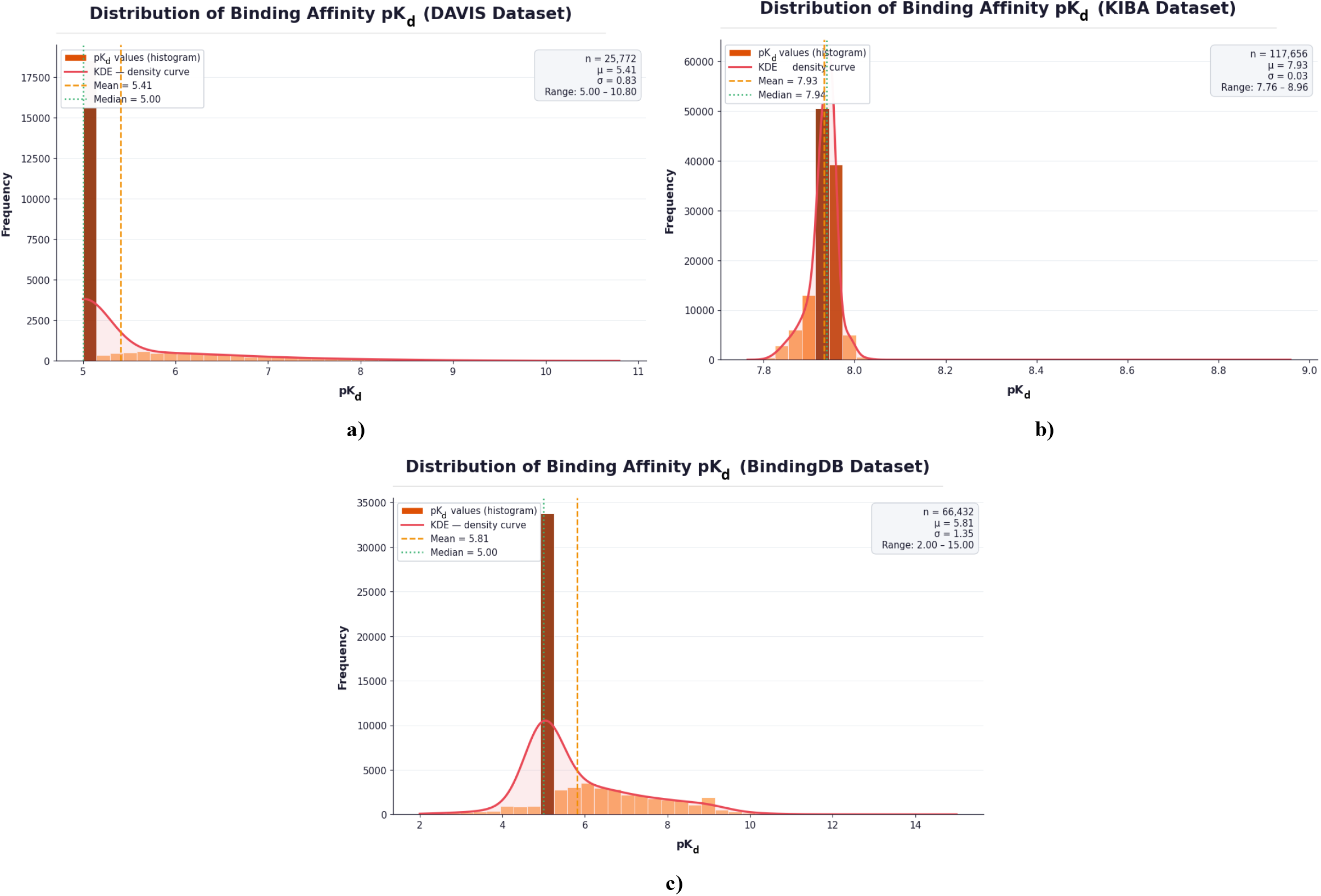
Distribution of p*K*_*d*_ values; a) DAVIS Dataset, b) KIBA Dataset, c) BDB Dataset

**Figure S4:**
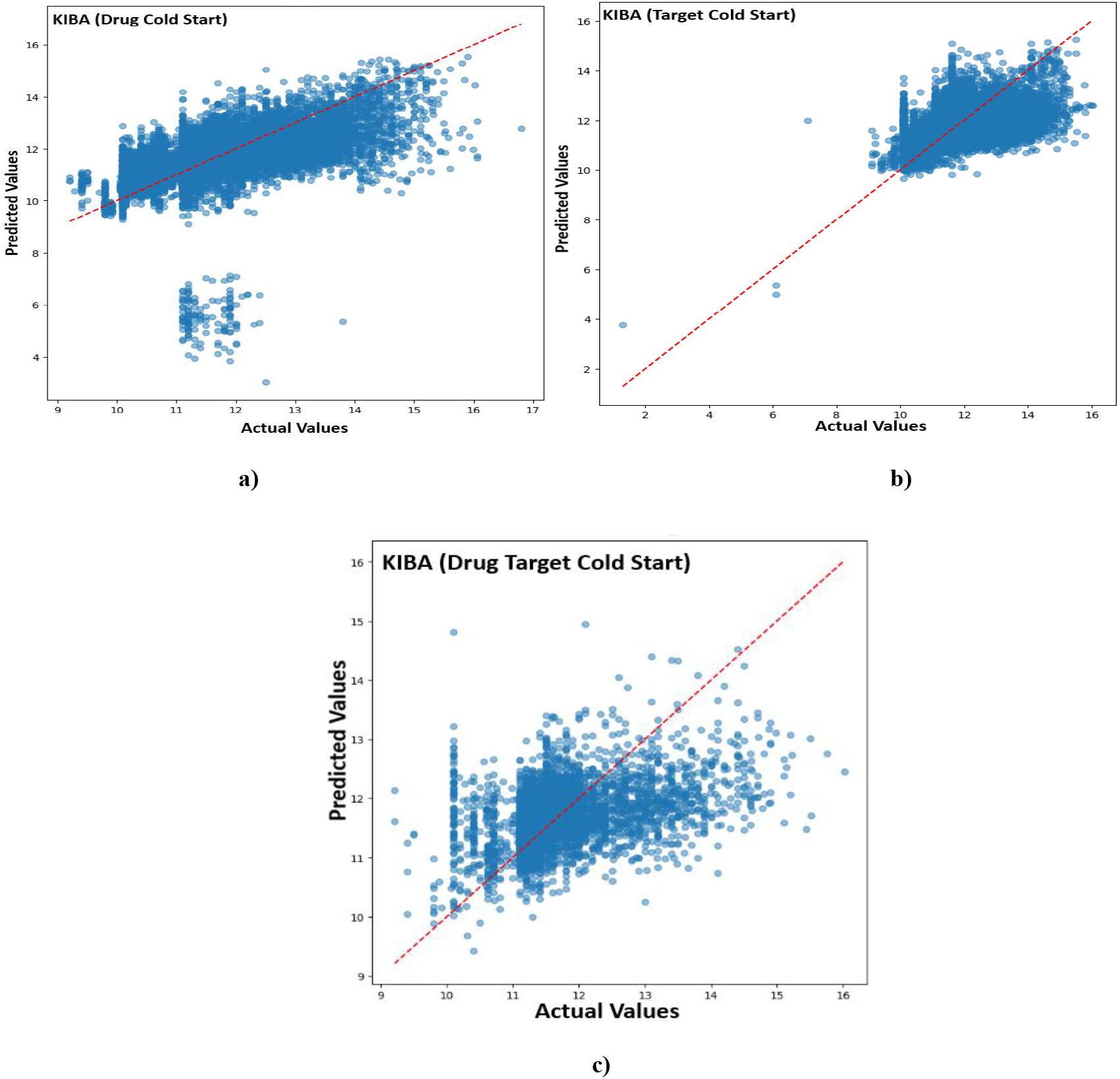
Scatter plots of the GTStrDTI - predicted *pK*_*d*_ against the experimentally measured *pK*_*d*_ on the KIBA dataset, a) Drug Cold Start, Target Cold Start, c) Drug Target Cold Start

**Figure S5:**
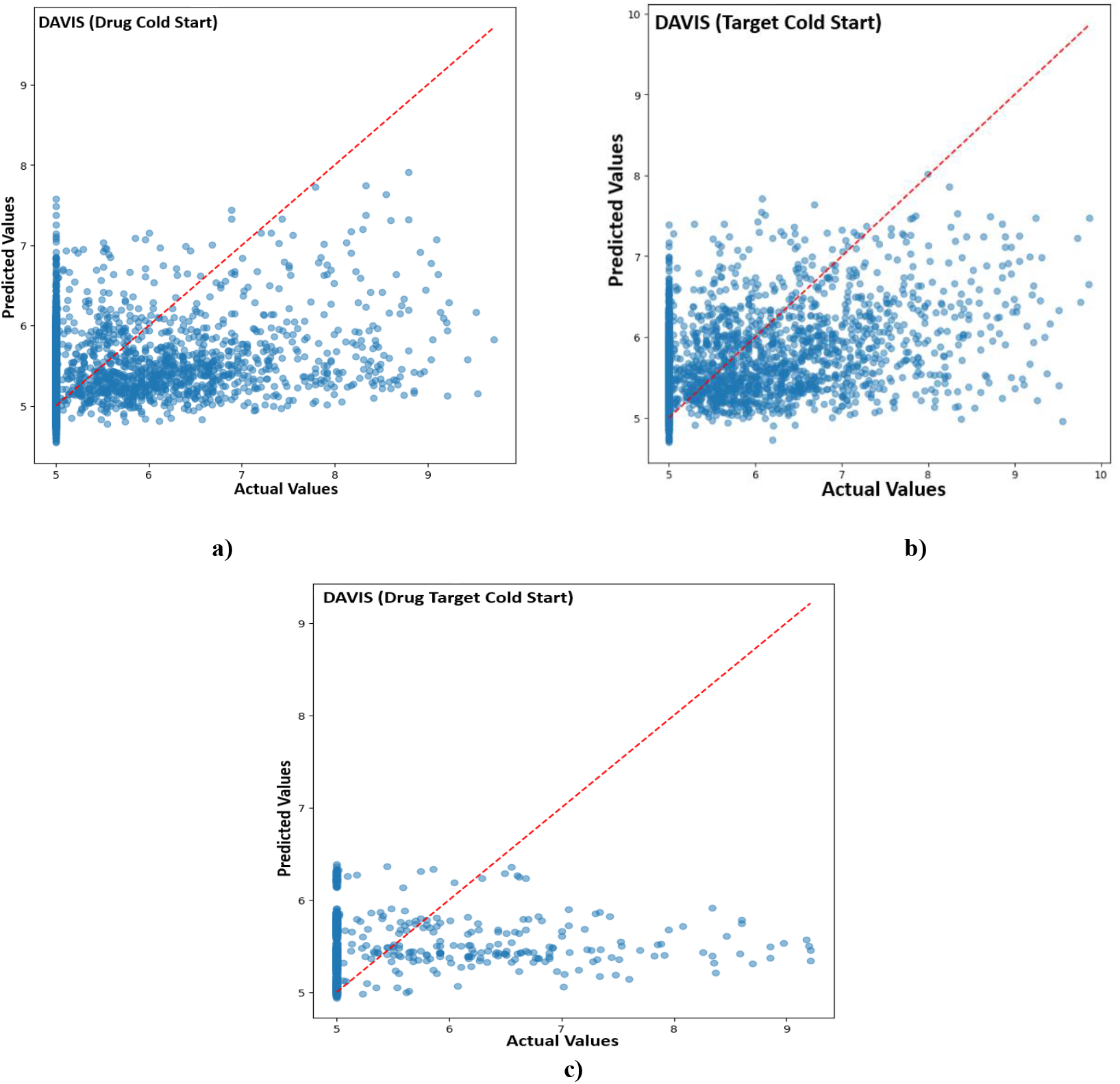
Scatter plots of the GTStrDTI - predicted *pK*_*d*_ against the experimentally measured *pK*_*d*_ on the DAVIS dataset, a) Drug Cold Start, b) Target Cold Start, c) Drug Target Cold Start

**Figure S6:**
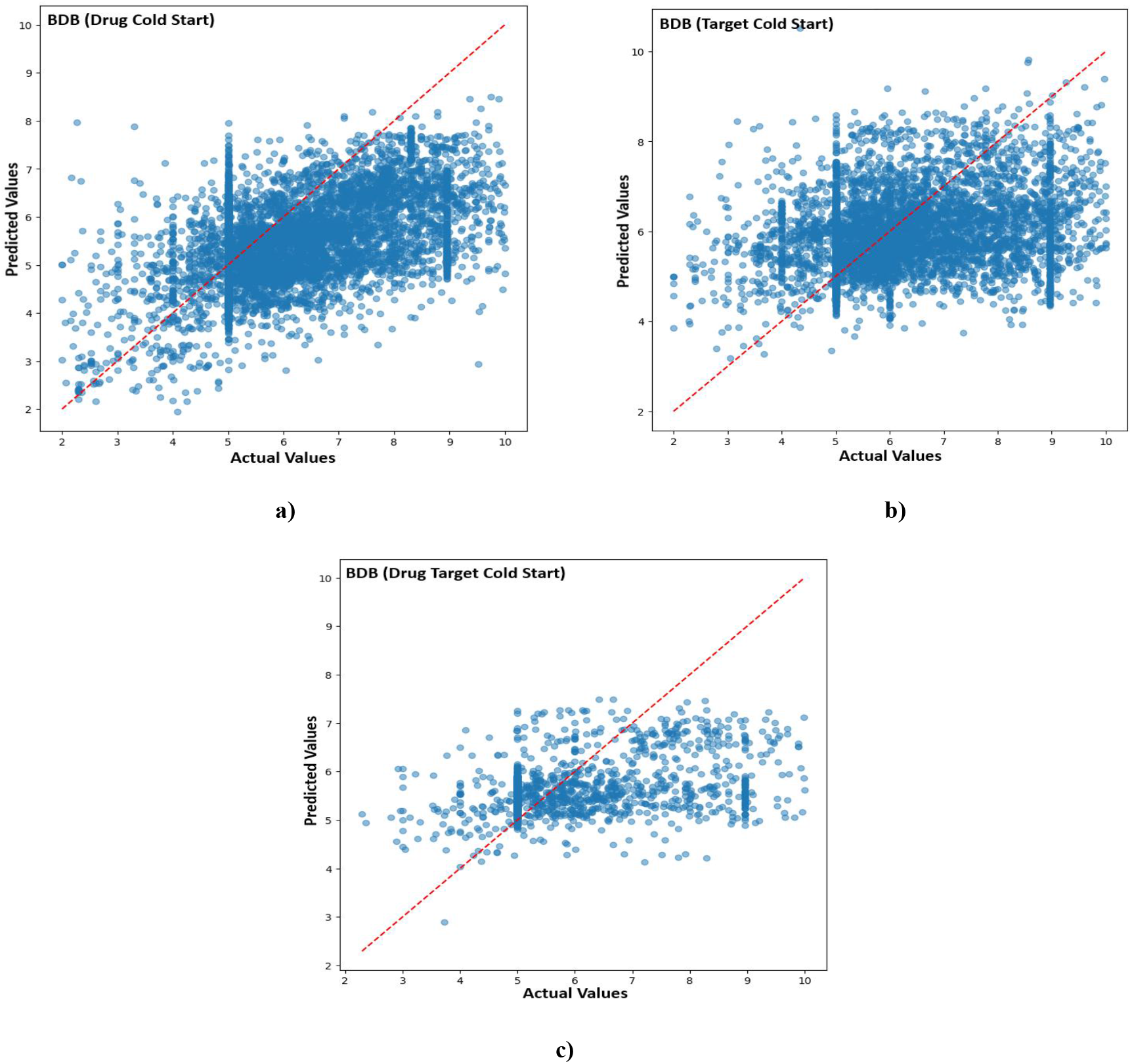
Scatter plots of the GTStrDTI - predicted *pK*_*d*_ against the experimentally measured *pK*_*d*_ on BDB data, a) Drug Cold Start, b) Target Cold Start, c) Drug Target Cold Start

**Table S1:** Data statistics for Cold Start Testing. The table summarizes the Distribution of training, validation, and test splits, reporting the number of drug target pairs along with the corresponding counts of unique drugs and unique targets in each split to evaluate model generalization under cold-start conditions.

| Datasets | Splits |  | Train | Validation | Test |
| --- | --- | --- | --- | --- | --- |
| **BDB** | Drug cold-start | Drug-Target Pairs | 30191 | 3933 | 8112 |
|  |  | Unique Drugs | 6921 | 989 | 1977 |
|  |  | Unique Target | 1028 | 684 | 792 |
|  | Target cold-start | Drug-Target Pairs | 29733 | 3891 | 8612 |
|  |  | Unique Drugs | 7552 | 1356 | 2492 |
|  |  | Unique Target | 761 | 109 | 218 |
|  | Drug-target cold -start | Drug-Target Pairs | 20184 | 503 | 1603 |
|  |  | Unique Drugs | 5207 | 160 | 468 |
|  |  | Unique Target | 716 | 74 | 164 |
| **DAVIS** | Drug cold-start | Drug-Target Pairs | 17813 | 2653 | 5306 |
|  |  | Unique Drugs | 47 | 7 | 14 |
|  |  | Unique Target | 379 | 379 | 379 |
|  | Target cold-start | Drug-Target Pairs | 18020 | 2584 | 5168 |
|  |  | Unique Drugs | 68 | 68 | 68 |
|  |  | Unique Target | 265 | 38 | 76 |
|  | Drug-target cold -start | Drug-Target Pairs | 12455 | 266 | 1064 |
|  |  | Unique Drugs | 47 | 7 | 14 |
|  |  | Unique Target | 265 | 38 | 76 |
| **KIBA** | Drug cold-start | Drug-Target Pairs | 84695 | 11254 | 21708 |
|  |  | Unique Drugs | 1447 | 207 | 414 |
|  |  | Unique Target | 229 | 224 | 228 |
|  | Target cold-start | Drug-Target Pairs | 80954 | 13943 | 22760 |
|  |  | Unique Drugs | 2068 | 1835 | 2058 |
|  |  | Unique Target | 160 | 23 | 46 |
|  | Drug-target cold -start | Drug-Target Pairs | 58271 | 1318 | 4189 |
|  |  | Unique Drugs | 1447 | 178 | 411 |
|  |  | Unique Target | 160 | 22 | 46 |

## REFERENCES

[1] Vajda, S., & Guarnieri, F. (2006). Characterization of protein-ligand interaction sites using experimental and computational methods. Current Opinion in Drug Discovery and Development, 9(3), 354. https://pubmed.ncbi.nlm.nih.gov/16729732/

[2] Kudari, Z. D., Kaira, V. S., Bhat, R., Raghavendra, V., Saxena, A., Jaiganesh, G., & Raghavendra, V. Advancing Protein-Ligand Binding Affinity Prediction: A Survey on Artificial Intelligence and Machine Learning Techniques. DOI: 10.36227/techrxiv.176344155.52503175/v1

[3] Rudrapal, M., & Chetia, D. (2020). Virtual screening, molecular docking and QSAR studies in drug discovery and development programme. Journal of drug delivery and therapeutics, 10(4), 225–233. 10.22270/jddt.v10i4.4218

[4] Huang, S. Y., & Zou, X. (2010). Advances and challenges in protein-ligand docking. International journal of molecular sciences, 11(8), 3016–3034. 10.3390/ijms11083016

[5] de Ruiter, A., & Oostenbrink, C. (2011). Free energy calculations of protein–ligand interactions. Current opinion in chemical biology, 15(4), 547–552. PMID: 21684797, DOI: 10.1016/j.cbpa.2011.05.021

[6] Khanna, V., & Ranganathan, S. (2022). Chemogenomics Approach to Computer Aided Drug Discovery. In Post-genomic Approaches in Drug and Vaccine Development (pp. 93–113). River Publishers. DOI:10.1201/9781003339090-5

[7] Ashish, V. (2017). Attention is all you need. Advances in neural information processing systems, 30, I. 10.48550/arXiv.1706.03762

[8] Friesner, R. A., Banks, J. L., Murphy, R. B., Halgren, T. A., Klicic, J. J., Mainz, D. T., … & Shenkin, P. S. (2004). Glide: a new approach for rapid, accurate docking and scoring. 1. Method and assessment of docking accuracy. Journal of medicinal chemistry, 47(7), 1739–1749. DOI: 10.1021/jm0306430

[9] Trott, O., & Olson, A. J. (2010). AutoDock Vina: improving the speed and accuracy of docking with a new scoring function, efficient optimization, and multithreading. Journal of computational chemistry, 31(2), 455–461. DOI: 10.1002/jcc.21334

[10] Ballester, P. J., & Mitchell, J. B. (2010). A machine learning approach to predicting protein–ligand binding affinity with applications to molecular docking. Bioinformatics, 26(9), 1169–1175. 10.1093/bioinformatics/btq112

[11] Li, G. B., Yang, L. L., Wang, W. J., Li, L. L., & Yang, S. Y. (2013). ID-Score: a new empirical scoring function based on a comprehensive set of descriptors related to protein– ligand interactions. Journal of chemical information and modeling, 53(3), 592–600. DOI: 10.1021/ci300493w

[12] Zilian, David, and Christoph A. Sotriffer. “Sfcscore rf: a random forest-based scoring function for improved affinity prediction of protein–ligand complexes.” Journal of chemical information and modeling 53.8 (2013): 1923–1933. DOI: 10.1021/ci400120b

[13] Kaira, V. S., Kudari, Z. D., Bhat, R., Saxena, A., & Jaiganesh, G. (2025). Deep Learning in Drug Discovery: Recent Advances, Applications and Future Directions. Authorea Preprints. DOI: 10.36227/techrxiv.176369451.19187650/v1

[14] Öztürk, H., Özgür, A., & Ozkirimli, E. (2018). DeepDTA: deep drug–target binding affinity prediction. Bioinformatics, 34(17), i821–i829. 10.1093/bioinformatics/bty593

[15] Suviriyapaisal, N., & Wichadakul, D. (2023). iEdgeDTA: integrated edge information and 1D graph convolutional neural networks for binding affinity prediction. RSC advances, 13(36), 25218–25228. 10.1039/D3RA03796G

[16] Jiménez, J., Skalic, M., Martinez-Rosell, G., & De Fabritiis, G. (2018). K deep: protein– ligand absolute binding affinity prediction via 3d-convolutional neural networks. Journal of chemical information and modeling, 58(2), 287–296. 10.1021/acs.jcim.7b00650

[17] Nguyen, Thin, et al. “GraphDTA: predicting drug–target binding affinity with graph neural networks.” Bioinformatics 37.8 (2021): 1140–1147. DOI: 10.1093/bioinformatics/btaa921

[18] Singh, Amritpal. “GraphPrint: extracting features from 3D protein structure for drug target affinity prediction.” arXiv preprint arXiv:2407.10452 (2024). 10.48550/arXiv.2407.10452

[19] Quan, L., Wu, J., Jiang, Y., Pan, D., & Qiang, L. (2025). DTA-GTOmega: enhancing drug-target binding affinity prediction with graph transformers using OmegaFold protein structures. Journal of Molecular Biology, 437(6), 168843. 10.1016/j.jmb.2024.168843

[20] Kyro, G. W., Smaldone, A. M., Shee, Y., Xu, C., & Batista, V. S. (2025). T-ALPHA: A hierarchical Transformer-Based deep neural network for Protein–Ligand binding affinity prediction with Uncertainty-Aware Self-Learning for Protein-Specific alignment. Journal of Chemical Information and Modeling, 65(5), 2395–2415. 10.1021/acs.jcim.4c02332

[21] Wu, Hongjie, et al. “AttentionMGT-DTA: A multimodal drug-target affinity prediction using graph transformer and attention mechanism.” Neural Networks 169 (2024): 623–636. 10.1016/j.neunet.2023.11.018

[22] Scarselli, F., Gori, M., Tsoi, A. C., Hagenbuchner, M., & Monfardini, G. (2008). The graph neural network model. IEEE transactions on neural networks, 20(1), 61–80. DOI: 10.1109/TNN.2008.2005605

[23] Landrum, G., Tosco, P., Kelley, B., Rodriguez, R., Cosgrove, D., Vianello, R., … & Monat, J. (2025). rdkit/rdkit: 2025_03_1 (Q1 2025) Release. Zenodo. DOI 10.5281/zenodo.15115844

[24] Oldfield, T. J., & Hubbard, R. E. (1994). Analysis of Cα geometry in protein structures. Proteins: Structure, Function, and Bioinformatics, 18(4), 324–337. 10.1002/prot.340180404

[25] Momin, Y., & Beloshe, V. (2025). Pharmacophore modeling in drug design. In Advances in Pharmacology (Vol. 103, pp. 313–324). Academic Press. 10.1016/bs.apha.2025.01.010

[26] Lee, J., Lee, I., & Kang, J. (2019, May). Self-attention graph pooling. In International conference on machine learning (pp. 3734–3743). pmlr. 10.48550/arXiv.1904.08082

[27] Clevert, D. A., Unterthiner, T., & Hochreiter, S. (2015). Fast and accurate deep network learning by exponential linear units (elus). arXiv preprint arXiv:1511.07289, 4(5), 11. 10.48550/arXiv.1511.07289

[28] Ba, J. L., Kiros, J. R., & Hinton, G. E. (2016). Layer normalization. arXiv preprint arXiv:1607.06450. 10.48550/arXiv.1607.06450

[29] Lu, J., Yang, J., Batra, D., & Parikh, D. (2016). Hierarchical question-image co-attention for visual question answering. Advances in neural information processing systems, 29. 10.48550/arXiv.1606.00061

[30] Nickel, M., & Tresp, V. (2013, September). Tensor factorization for multi-relational learning. In Joint European Conference on Machine Learning and Knowledge Discovery in Databases (pp. 617–621). Berlin, Heidelberg: Springer Berlin Heidelberg. 10.1007/978-3-642-40994-3_40

[31] Tang, J., Szwajda, A., Shakyawar, S., Xu, T., Hintsanen, P., Wennerberg, K., & Aittokallio, T. (2014). Making sense of large-scale kinase inhibitor bioactivity data sets: a comparative and integrative analysis. Journal of chemical information and modeling, 54(3), 735–743. 10.1021/ci400709d

[32] Davis, Mindy I., et al. “Comprehensive analysis of kinase inhibitor selectivity.” Nature biotechnology 29.11 (2011): 1046–1051. DOI: 10.1038/nbt.1990

[33] Liu, T., Lin, Y., Wen, X., Jorissen, R. N., & Gilson, M. K. (2007). BindingDB: a web-accessible database of experimentally determined protein–ligand binding affinities. Nucleic acids research, 35(suppl_1), D198–D201. 10.1093/nar/gkl999

[34] Barbet, J., & Huclier-Markai, S. (2019). Equilibrium, affinity, dissociation constants, IC5O: Facts and fantasies. Pharmaceutical Statistics, 18(5), 513–525. 10.1002/pst.1943

[35] Huang, K., Fu, T., Gao, W., Zhao, Y., Roohani, Y., Leskovec, J., & Zitnik, M. (2021). Therapeutics data commons: Machine learning datasets and tasks for drug discovery and development. arXiv:2102.09548. 10.48550/arXiv.2102.09548

[36] Steck, H., Krishnapuram, B., Dehing-Oberije, C., Lambin, P., & Raykar, V. C. (2007). On ranking in survival analysis: Bounds on the concordance index. Advances in neural information processing systems, 20. https://www.researchgate.net/publication/221618691_On_Ranking_in_Survival_Analysis_Bounds_on_the_Concordance_Index

